# Stochastic RNA editing of the Complexin C-terminus within single neurons regulates neurotransmitter release

**DOI:** 10.1101/2023.05.30.542887

**Authors:** Elizabeth A. Brija, Zhuo Guan, Suresh K. Jetti, J. Troy Littleton

## Abstract

Neurotransmitter release requires assembly of the SNARE complex fusion machinery, with multiple SNARE-binding proteins regulating this process to control when and where synaptic vesicle fusion occurs. Complexin (Cpx) controls spontaneous and evoked neurotransmitter release by modulating SNARE complex zippering. Although the central SNARE-binding helix is essential, post-translational modifications to Cpx’s C-terminal membrane-binding amphipathic helix modulate its activity. Here we demonstrate that RNA editing of the Cpx C-terminus regulates its ability to clamp SNARE-mediated fusion and alters presynaptic output. RNA editing of Cpx within single neurons is stochastic, generating up to eight edit variants that fine-tune neurotransmitter release by changing the subcellular localization and clamping properties of the protein. Similar editing rules for other synaptic genes were observed, indicating stochastic editing at single adenosines and across multiple mRNAs can generate unique synaptic proteomes within the same population of neurons to fine-tune presynaptic output.

## Introduction

Neuronal communication is initiated by Ca^2+^-evoked fusion of synaptic vesicles (SVs) in response to action potentials. Single SVs also fuse spontaneously to generate mini events. Complexin (Cpx) and the Ca^2+^ sensor Synaptotagmin 1 (Syt1) bind and regulate SNARE complexes to control whether SVs fuse spontaneously or through the evoked pathway^1–3^. Cpx can arrest zippering of the SNARE complex at the SV/plasma membrane interface to maintain SVs in a fusion-ready state and allow Ca^2+^-bound Syt1 to rapidly trigger release^4–6^. Indeed, invertebrate Cpxs act as “fusion clamps” to reduce spontaneous release in the absence of Ca^2+^^7–10^. Changes in Cpx activity alters spontaneous release rates and regulates structural and functional synaptic plasticity^7,11–14^. The Cpx C-terminus has emerged as a key site for such regulatory control, as it encodes a conserved amphipathic helix that functions as a membrane curvature sensor to localize Cpx to SVs and concentrate its activity at release sites^15^.

In *Drosophila*, a single *cpx* gene produces two isoforms with different C-termini due to alternative splicing of exon 7^16^. The Cpx7A isoform has a conserved membrane-tethering prenylation CAAX box, while Cpx7B lacks this motif^17^. Prior studies demonstrated Cpx7B is regulated by PKA phosphorylation of a serine residue (S126) within this alternatively spliced C-terminus region. PKA phosphorylation of Cpx7B reduces its clamping function at neuromuscular junctions (NMJs), leading to elevated spontaneous release that triggers activity-induced structural plasticity^11^. The Cpx7A isoform lacks this serine residue and instead undergoes RNA editing by ADAR (adenosine deaminase acting on RNA) to generate multiple Cpx7A proteins with unique C-terminal sequences. A-to-I editing can recode pre-spliced mRNAs by deaminating target adenosines in double-stranded RNA structures induced from exon-intron complementary pairing, causing the resulting inosine base to be read as guanosine by the translation machinery^18^. RNA editing of Cpx7A can trigger changes at two resides, including an isoleucine (I125) to methionine substitution. In addition, editing of two adjacent downstream adenosines can change an asparagine (N130) to a glycine (N130G), aspartate (N130D), or serine (N130S) at a site near the phosphorylated S126 residue in Cpx7B^16,19^. Given Cpx7A is expressed at higher levels and is the dominant isoform in the *Drosophila* nervous system^16^, RNA editing at this site represents an attractive mechanism for regulating neurotransmitter release and structural plasticity across a larger population of neurons.

Here we used CRISPR and transgenic rescue to assay the role of Cpx splicing and RNA editing in neurotransmitter release. Although Cpx7A is expressed at higher levels, the two isoforms are largely redundant in their ability to support baseline synaptic transmission. Analysis of single-cell RNAseq data from individual motoneurons reveals multiple Cpx7A RNA editing variants can be simultaneously expressed, indicating RNA editing does not act in an “all-or-none” fashion. The most prominent edit variant (Cpx7A^I125M,N130S^) can be phosphorylated by Casein kinase 2 (CK2). Transgenic rescue of *cpx* null mutants with Cpx7A^I125M,N130S^ demonstrates RNA editing alters the protein’s subcellular localization and reduces its ability to clamp spontaneous SV fusion, leading to synaptic overgrowth. Rescue with both Cpx7A^I125M,N130S^ and unedited Cpx7A^I125,N130^ indicates the N130S variant acts in a dominant fashion, consistent with a model where multiple Cpxs engage assembling SNARE complexes during SV fusion^20^. Such a mechanism would allow edited and unedited Cpx proteins to assert independent effects in a combinatorial fashion to control fusion dynamics. Together, these data indicate stochastic RNA editing of the Cpx C-terminus can set distinct spontaneous release rates in individual *Drosophila* neurons to regulate presynaptic output.

## Results

### Alternative splicing and RNA editing of *Drosophila complexin* generates divergent C-terminal sequences within a conserved amphipathic helical domain

In contrast to four Cpx homologs in mammals, a single *cpx* gene is present in *Drosophila*. *Drosophila cpx* undergoes alternative splicing of exon 7 to generate two unique isoforms, Cpx7A and Cpx7B, that differ in their last ~20 amino acids (Figure 1A, B). Although the encoded exon 7 sequences are not similar at the amino acid level, the C-terminus of both splice isoforms encode a membrane-binding amphipathic helix (Figure 1C, D) that is conserved across invertebrate and vertebrate Cpx homologs^15^. Cpx7A is the more abundant isoform and contains a C-terminal CAAX box that undergoes prenylation^16^, a post-translational lipid attachment that helps localize this variant and mammalian CPX3 and CPX4 within synapses^12,17,21^. The less abundant Cpx7B lacks a prenylation motif, similar to mammalian CPX1 and CPX2. We previously demonstrated the Cpx7B C-terminus is phosphorylated by PKA at residue S126 in an activity-dependent manner, leading to reduced SV clamping and enhanced spontaneous release and synaptic growth^11^.

**Figure 1.**
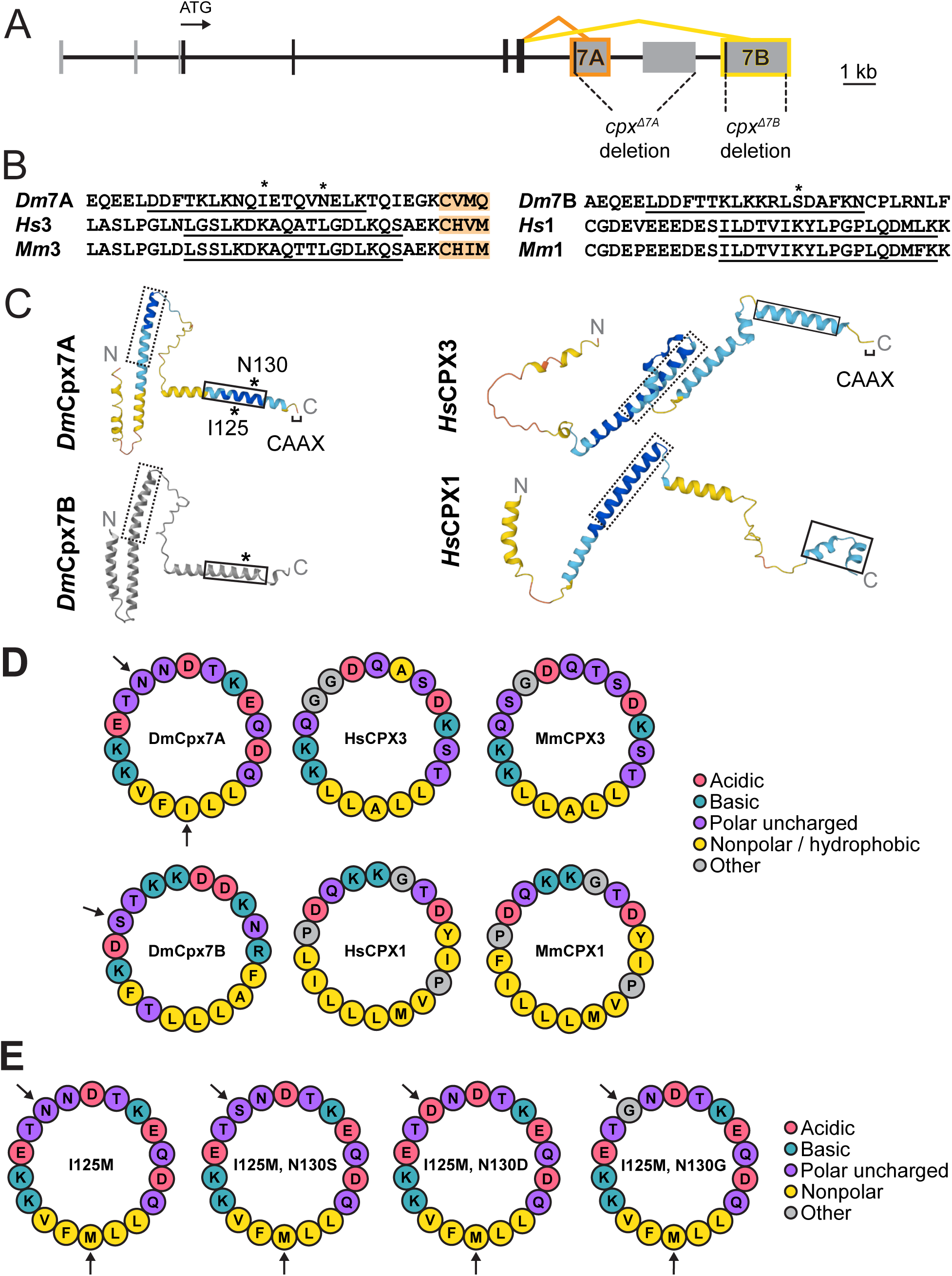
Alternative splicing and RNA editing generate diversity in the conserved Complexin C-terminal amphipathic helix. (**A**) Diagram of the *Drosophila cpx* genomic locus with protein-coding exons indicated by black boxes and noncoding exons with gray. The ATG start codon is noted, together with the 7A (orange) and 7B (yellow) alternative splicing events. The location of CRISPR-generated *cpx* deletions removing 7A (*cpx^Δ7A^*) or 7B (*cpx^Δ7B^*) are also shown. (**B**) Alignment of Cpx C-termini from several species (Dm – *Drosophila melanogaster*, Hs – *Homo sapiens*, Mm – *Mus musculus*) highlights the two subfamilies that contain or lack a CAAX prenylation motif (orange). The amphipathic helix region is underlined, with asterisks denoting residues modified by RNA editing (I125, N130) in *Dm*Cpx7A or phosphorylation (S126) in *Dm*Cpx7B. (**C**) AlphaFold predictions of Cpx structure for homologs with and without the CAAX prenylation motif. The dashed line box denotes the SNARE-binding central helix and the solid line box highlights the C-terminal amphipathic helix. Cpx7A RNA editing sites and the Cpx7B phosphorylation site are denoted with *. AlphaFold per-residue confidence scores (pLDDT) are color-coded: blue = very high (pLDDT > 90), cyan = confident (90 > pLDDT > 70), yellow = low (70 > pLDDT > 50), orange = very low (pLDDT < 50). Cpx7B was generated using a simplified AlphaFold version without confidence scores and visualized with iCn3D^72^. (**D**) HELIQUEST predictions of the Cpx C-terminal amphipathic helix show the conserved hydrophilic and hydrophobic faces, with amino acid properties noted in the legend on the right. Arrows indicate the Cpx7A edit sites and the Cpx7B phosphorylation site. (**E**) Amphipathic helix models for non-edited (left) and edited (right) Cpx7A proteins.

Given the regulatory role of Cpx7B phosphorylation, it was surprising the more abundant Cpx7A lacks this PKA phosphorylation site. Unlike Cpx7B, Cpx7A is subject to RNA editing via ADAR at three adenosine residues within the mRNA sequence of exon 7A^16,19^. One edit site generates a isoleucine (I) to methionine (M) conserved substitution at amino acid 125 that is not predicted to alter protein function, but induces mRNA conformational changes in exon-intron base pairing that facilitates editing of two downstream adenosine residues^16^. At this downstream site, the unedited AAT codon encodes an asparagine (N) at amino acid 130. Editing of both residues (AAT to GGT) produces a glycine (N130G), while editing of only the 1^st^ base (AAT to GAT) generates an aspartic acid (N130D) and editing of the 2^nd^ base (AAT to AGT) produces a serine (N130S) (Figure 1E). Given RNA editing generates a potentially phospho-competent Cpx7A^N130S^, a phospho-mimetic Cpx7A^N130D^ and a phospho-incompetent Cpx7A^N130G^, these editing changes could control Cpx7A function. To begin testing this hypothesis, a structural comparison of Cpx7A and Cpx7B with their mammalian homologs was performed using AlphaFold^22,23^. AlphaFold predictions indicate each Cpx homolog contains a conserved SNARE-binding central helix and a C-terminal helical domain (Figure 1C).

Given the C-terminal amphipathic helix of Cpx detects membrane curvature and binds SVs^24–27^, helical wheel models were generated with HELIQUEST^28^ to examine hydrophilic and hydrophobic faces of the helix in relation to the Cpx7A N130 editing site and the Cpx7B S126 PKA phosphorylation site (Figure 1D). Both splice isoforms have phospho-competent serine and/or threonine residues in similar positions on the hydrophilic face, including the Cpx7B S126 residue. Mammalian CPXs also contain phospho-competent residues on this hydrophilic surface (Figure 1D), suggesting phosphorylation of this region may represent a conserved mechanism for modulating Cpx activity. The I125M edit resides on the hydrophobic face of the helix and does not alter the hydrophobic nature of this region (Figure 1E). In contrast, the Cpx7A N130S edit adds another phospho-competent residue to match the paired S/T residues where Cpx7B S126 resides (Figure 1E). The N130D edit adds another negative charge to the hydrophilic face, generating a helix with a negative charge at nearly every other amino acid on this surface. The N130G edit inserts a glycine residue that exactly matches glycine residues found at the same site in mammalian CPX3, consistent with a functional impact for this editing event as well. We conclude that RNA editing alters the hydrophilic face of the Cpx7A C-terminal amphipathic helix to generate variants that more closely resemble Cpx7B or mammalian CPXs, suggesting RNA editing may alter the properties of the helix or its potential for phosphorylation.

### Characterizing the functional significance of alternative splicing of Cpx exon 7

Before characterizing the impact of RNA editing on Cpx7A function, we first examined the role of Cpx alternative splicing to gain greater insight into the endogenous roles of Cpx7A and Cpx7B. In *cpx* null mutants that lack both isoforms (*cpx^SH^*^1^), spontaneous mini frequency is dramatically elevated (>50-fold), evoked release is decreased, and larval NMJ synaptic growth is enhanced^7^. Overexpression of either Cpx7A or Cpx7B in *cpx^SH1^* partially rescues these phenotypes, with Cpx7A having more robust clamping properties and Cpx7B over-rescuing evoked release^16^. Although both variants support aspects of Cpx function when overexpressed, endogenous Cpx7A mRNA is >100-fold more abundant than Cpx7B in *Drosophila* adults and larvae based on quantitative RT-PCR^16^. As such, Cpx7A is hypothesized to play a more critical role in synaptic transmission. To directly test the endogenous function of the two splicing isoforms, CRISPR mutants disrupting 7A (*cpx^Δ7A^*) or 7B (*cpx^Δ7B^*) were generated by introducing an early stop codon at the beginning of exon 7A or 7B, respectively. Western analysis of adult head lysates with a Cpx antibody that recognizes both variants showed an 87% reduction in overall Cpx expression in *cpx^Δ7A^* (*p* ˂ 0.0001) and a milder 10% reduction in *cpx^Δ7B^*(*p* = 0.9398), consistent with Cpx7A being the predominant isoform (Figure 2A). Immunostaining for Cpx at 3^rd^ instar larval NMJs showed a similar effect (Figure 2B, C), with a >85% reduction in Cpx at synapses in *cpx^Δ7A^* mutants (*p* ˂ 0.0001) and no detectable decrease in *cpx^Δ7B^* mutants (*p* = 0.9567).

**Figure 2.**
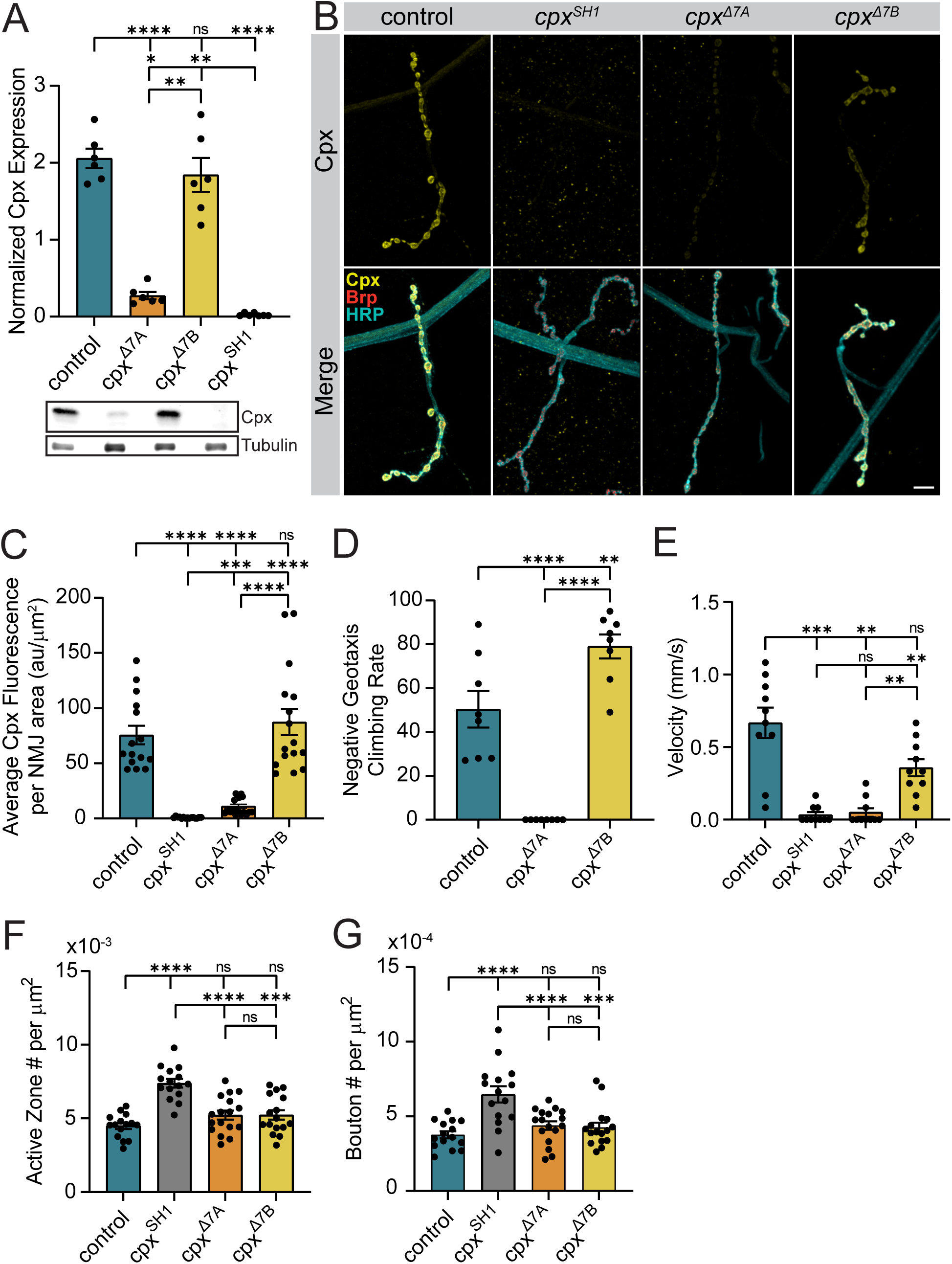
Morphological and behavioral phenotypes in CRISPR-generated splicing mutants lacking Cpx exon 7A or 7B. **(A)** Quantification and representative western blot of Cpx levels from adult head extracts normalized to the loading control (anti-Tubulin) for the indicated genotypes (control (*white*), *cpx^Δ7A^*, *cpx^Δ7B^*, and *cpx^SH1^*). **(B)** Immunostaining of 3^rd^ instar larval muscle 4 NMJs at segment A3 for the indicated genotypes with antibodies against Cpx (yellow, upper panels), Brp (red) and anti-HRP (cyan) show the large decrease in total Cpx levels in *cpx^Δ7A^* mutants lacking the predominant Cpx7A isoform. Scale bar = 10 µm. **(C)** Quantification of total Cpx fluorescence within the HRP-positive area at muscle 4 NMJs for the indicated genotypes (au = arbitrary units). **(D)** Quantification of climbing rate in negative geotaxis assays for adult males of the indicated genotypes demonstrates severe motor deficits in *cpx^Δ7A^* mutants compared to *cpx^Δ7B^*. Each point represents the average climbing rate for a cohort of 10 males. **(E)** Quantification of 3^rd^ instar larval crawling velocity for the indicated genotypes. **(F)** Quantification of mean AZ number per muscle area at muscle 4 NMJs of the indicated genotypes. **(G)** Quantification of mean synaptic bouton number per muscle area at muscle 4 NMJs of the indicated genotypes. Data are shown as mean ± SEM; \**p* ˂ 0.05, \*\**p* ˂ 0.01, \*\*\**p* ˂ 0.001, \*\*\*\**p* ˂ 0.0001, ns = not significant.

To assay functional requirements for the splice variants *in vivo*, adult and larval motor behavior were examined in *cpx^Δ7A^* and *cpx^Δ7B^* and compared to control and *cpx^SH1^* nulls. Complete loss of Cpx in *cpx^SH1^* severely disrupts behavior and reduces viability, with the few escaper adults that emerge from the pupal case displaying a profound loss of motor control and an inability to walk in a coordinated manner. Loss of the predominant Cpx7A isoform also strongly disrupted motor behavior, though not as severely as *cpx^SH1^*. In contrast to *cpx^SH1^* where homozygous adults are rarely observed, *cpx^Δ7A^* could be maintained as a homozygous stock, indicating the remaining endogenous Cpx7B can improve viability and fertility compared to animals lacking both isoforms. However, *cpx^Δ7A^* adults showed a complete inability to climb in a negative geotaxis assay (Figure 2D). In contrast, *cpx^Δ7B^* adults were fully viable and did not display obvious motor defects. In addition, *cpx^Δ7B^* adults were moderately hyperactive in climbing compared to controls (control: 50.4 ± 8.3% pass rate, *n* = 8 cohorts of ten flies; *cpx^Δ7A^*: 0 ± 0% pass rate, *n* = 8 cohorts of ten flies, *p* ˂ 0.0001 to control; *cpx^Δ7B^*: 79 ± 5.5% pass rate, *n* = 8 cohorts of ten flies, *p* = 0.0055 to control, *p* ˂ 0.0001 to *cpx^Δ7A^*). To examine larval locomotion, crawling velocity of 3^rd^ instar larvae was assayed in *cpx^Δ7A^* and *cpx^Δ7B^* mutants (Figure 2E). Similar to *cpx^SH1^*, *cpx^Δ7A^* larvae displayed a large reduction in crawling velocity (*p* = 0.0011). In contrast, *cpx^Δ7B^* mutants showed only a mild decrease in velocity (*p* = 0.1155). Together, these data indicate Cpx7A has a more prominent role in supporting larval and adult motor behavior.

To examine synaptic morphology and neurotransmitter release in *cpx^Δ7A^*and *cpx^Δ7B^* mutants, the well-characterized 3^rd^ instar larval glutamatergic NMJ preparation was used^29^. Synaptic bouton and active zone (AZ) number were quantified by immunostaining for neuronal membranes (anti-HRP) and the AZ protein Bruchpilot (Brp) (Figure 2B, F, G). In contrast to the large increase in bouton (72%, *p* = 0.0012) and AZ (65%, *p* < 0.0001) number in *cpx^SH1^*, synaptic growth was largely unaffected in *cpx^Δ7A^* (AZ#: *p* = 0.2809, bouton#: *p* = 0.4576) and *cpx^Δ7B^* (AZ#: *p* = 0.3121, bouton#: *p* = 0.7781) mutants. Despite the differences in overall levels of the two splice isoforms at NMJs (Figure 2B), *cpx^Δ7A^* mutants had similar NMJ morphology to *cpx^Δ7B^* (*p* = 0.9999). We conclude that Cpx7A and Cpx7B are functionally redundant in their ability to regulate synaptic growth, with Cpx7B having a more important role than expected based on its lower expression level.

To assay synaptic function, two-electrode voltage-clamp (TEVC) was used to measure evoked and spontaneous neurotransmitter release at 3^rd^ instar larval muscle 6 in abdominal segment A3. In contrast to the dramatic increase in spontaneous release rate observed in *cpx^SH1^* (>55-fold), endogenously expressed Cpx7A or Cpx7B was able to substantially lower mini frequency (Figure 3A, B). The presence of only Cpx7A in the *cpx^Δ7B^* mutant returned spontaneous release rates to control levels (*p* = 0.2641). The residual Cpx7B in *cpx^Δ7A^* mutants was not able to fully clamp spontaneous fusion, with a 2-fold increase compared to controls (*p* ˂ 0.0001). In contrast to spontaneous release, the presence of only one of the two splice isoforms was not as effective in driving normal levels of evoked fusion (Figure 3C, D). A 71% reduction in the peak amplitude of the evoked excitatory junctional current (eEJC) was observed in *cpx^SH1^* compared to control (*p* ˂ 0.0001). Although less severe, *cpx^Δ7A^*displayed a 46% reduction (*p* ˂ 0.0001) and *cpx^Δ7B^* a 25% reduction (*p* = 0.0269) compared to controls. These data indicate Cpx7A and Cpx7B are both required for wildtype evoked responses, with the higher levels of Cpx7A in *cpx^Δ7B^* mutants supporting more evoked fusion than the lower levels of Cpx7B in *cpx^Δ7A^*.

**Figure 3.**
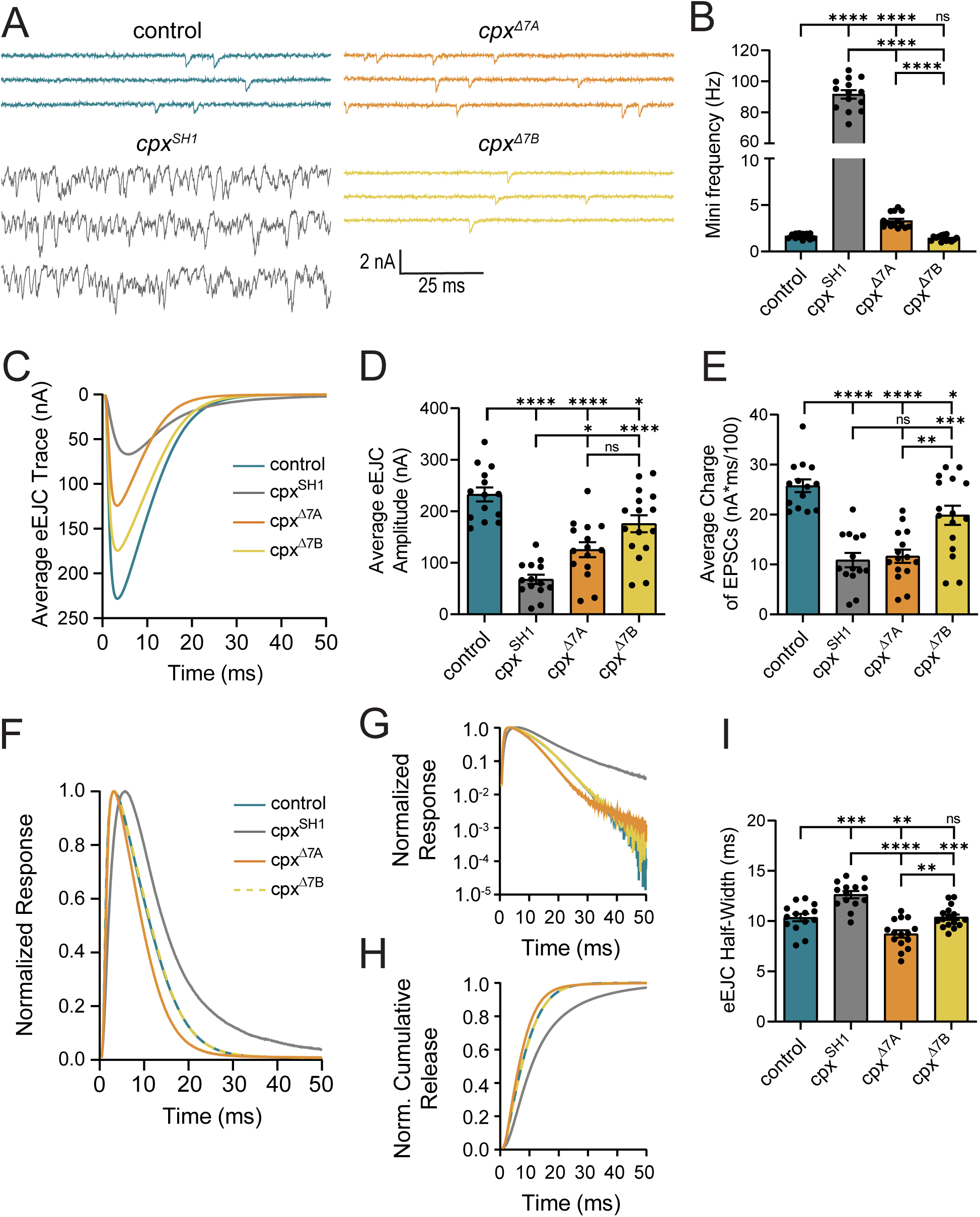
Electrophysiological analysis of synaptic function in mutants lacking Cpx7A or Cpx7B. **(A)** Representative postsynaptic current recordings of spontaneous release at muscle 6 NMJs in control (blue), *cpx^SH1^* (gray), *cpx^Δ7A^* (orange), or *cpx^Δ7B^* (yellow) 3^rd^ instar larvae. This genotypic color scheme is maintained for all figure panels. **(B)** Quantification of average spontaneous release rate for the indicated genotypes. Note the y-axis gap between 10-60 Hz due to the extreme elevation of mini frequency in *cpx^SH1^* null mutants. **(C)** Average traces of evoked EJC responses for the indicated genotypes. **(D)** Quantification of average eEJC amplitude for the indicated genotypes. **(E)** Quantification of average evoked release charge obtained by measuring total release over time following single action potentials. **(F)** Average normalized evoked responses for each genotype. *cpx^Δ7B^* is depicted with dashed yellow line for ease of visualization. **(G)** Average normalized responses plotted on a semi-logarithmic graph to display release components for each genotype. Note the large increase in the slower asynchronous release component in *cpx^SH1^* null mutants (gray line). **(H)** Cumulative release normalized for the maximum response in 2.0 mM external Ca^2+^ for each genotype. Each trace was adjusted to a double exponential fit. *cpx^Δ7B^*is depicted with dashed yellow line for ease of visualization. **(I)** Average evoked EJC half-width change for each genotype. All recordings were performed in 2.0 mM external Ca^2+^ saline. Data are shown as mean ± SEM; \**p* ˂ 0.05, \*\**p* ˂ 0.01, \*\*\**p* ˂ 0.001, \*\*\*\**p* ˂ 0.0001, ns = not significant.

In addition to the total number of SVs that fuse during an action potential, Cpx also modulates SV release kinetics by promoting fast synchronous fusion and reducing the slower asynchronous pathway^4^. While peak eEJC amplitude primarily captures synchronous release, eEJC charge, half-width, and the time course of cumulative release provide kinetic insights into synchronous and asynchronous fusion. These evoked variables were compared across control, *cpx^Δ7A^*, *cpx^Δ7B^* and *cpx^SH1^* NMJs (Figure 3E-I). Like eEJC amplitude, release kinetics in *cpx^Δ7B^* mutants were more similar to controls. Loss of Cpx7A in the *cpx^Δ7A^* line resulted in reduced charge transfer (Figure 3E) and a mild increase in asynchronous release (Figure 3F, G), though less than the large amount of asynchronous fusion in *cpx* nulls. The onset of synchronous release was slightly enhanced in *cpx^Δ7A^* mutants (Figure 3G, H), consistent with prior data showing overexpression of Cpx7B enhanced the speed of onset of evoked fusion^4^. In summary, we conclude endogenous levels of either Cpx isoform are sufficient to clamp spontaneous SV fusion. The requirement for both isoforms to fully recapitulate evoked responses indicate they have both shared and independent roles in Ca^2+^-dependent SV release. Given the more severe behavioral defects observed in animals lacking Cpx7A, some neuronal subtypes are likely to be more highly reliant on this splice variant for supporting synaptic transmission, in contrast to motoneurons.

### Single-cell RNAseq reveals stochastic RNA editing of Cpx7A in larval motoneurons

Having analyzed functions for the two C-terminal splice isoforms of Cpx, we next examined RNA editing profiles for Cpx7A in larval motoneurons. Larval abdominal muscles are innervated by several populations of glutamatergic motoneurons in *Drosophila*, including tonic Type Ib and phasic Type Is subclasses^30,31^. Type Ib and Is motoneurons are the primary drivers of muscle contraction and display unique morphological and functional properties^32–37^. Given Ib and Is neurons have distinct presynaptic release output, we hypothesized RNA editing of Cpx7A could contribute to these differences if: (1) Cpx7A edit variants were differentially expressed between the two neuronal populations; and (2) editing of Cpx7A altered its function. To determine the abundance and diversity of Cpx7A edit variants (I125M, N130G, N130S, N130D), single-cell RNAseq datasets we previously generated from ~200 larval Ib and Is motoneurons were analyzed^33^. For these experiments, whole cell electrodes were used to collect cytosolic and nuclear content from individual Ib or Is neurons that expressed GFP using Gal4 drivers specific to each cell type^32^. High-resolution paired-end deep single-cell RNA sequencing was performed on RNA extracted from each individual motoneuron, generating ~4 million reads per cell and allowing identification of RNA editing diversity at single neuron resolution. Prior studies using mRNA obtained from pooled neuronal populations proposed RNA editing occurs in an “all-or-none” fashion at individual edit sites within a cell. For example, more than 99% of mammalian AMPA GluA2 receptor subunit transcripts undergo RNA editing at specific times during brain development^38,39^. Given our method provides individual neuron resolution of RNA editing, we could directly test the “all-or-none” model. Strikingly, individual larval motoneurons showed highly stochastic RNA editing of the three adenosine bases (position 375, 388 and 389) that are subject to possible editing in *Cpx* exon 7A. Editing rates for these adenosines ranged from 0 to 100% across the ~200 neurons, with 98% of Ib cells and 94% of Is cells showing some level of exon 7A editing (Figure 4A, B). For the adenosine 375 edit site, the average I125M (adenosine 375 to inosine 375) edited transcript level per neuron was 31.7 ± 2.4% in Ib (*n* = 95 cells) and 30.3 ± 2.4% in Is (*n* = 86 cells). Only 3% of Ib and 1% of Is neurons fully edited all Cpx mRNA to I125M. For all Cpx7A mRNA edit variants, the average Ib motoneuron expressed 53% unedited Cpx, 32% Cpx^I125M^, 0.0% Cpx^N130D^, 1.5% Cpx^N130S^, 0.3% Cpx^N130G^, 0.2% Cpx^I125M,N130D^, 9.5% Cpx^I125M,N130S^ and 3.7% Cpx^I125M,N130G^ (Figure 4A, B). The average Is motoneuron expressed a similar ratio of edited Cpx7A transcripts (Figure 4A, B). As such, differential RNA editing of Cpx7A is unlikely to drive the distinct release properties of Ib and Is motoneurons, though stochastic editing of Cpx7A could contribute to individual neuronal heterogeneity in presynaptic output. We conclude that RNA editing of Cpx exon 7A is stochastic across individual edit sites and generates similar diversity of edited isoforms in Is and Ib motoneurons.

**Figure 4.**
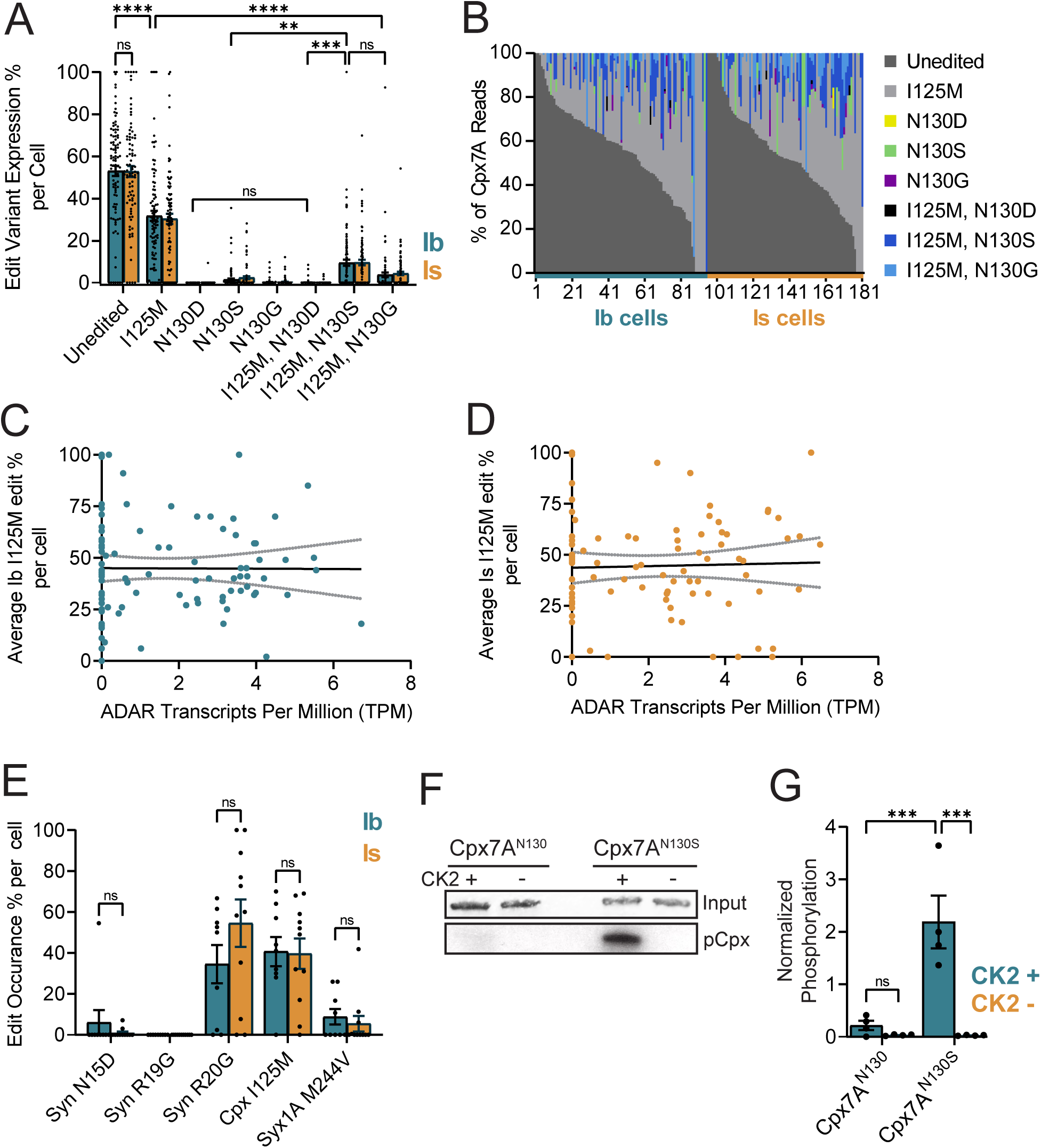
Stochastic expression of Cpx7A RNA editing variants in single neurons. **(A)** Quantification of unedited and edited Cpx7A mRNAs from single Ib (blue, *n* = 95 cells) and Is (orange, *n* = 86 cells) motoneuron RNAseq datasets. Each point represents the number of edit variant reads as percent of total Cpx reads in an individual neuron. **(B)** Sorted single-cell RNA editing profiles for Cpx7A across the sampled population of Ib and Is motoneurons. Each neuron is displayed as a stacked bar with corresponding edit and unedited read percentages that total 100% of *cpx* mRNA for that cell. Cells 1-95 are Ib motoneurons and cells 96-181 are Is motoneurons. Neurons are sorted by the largest unedited percent for both Ib and Is groups. (**C**) Comparison of ADAR transcripts per million (TPM) profile in Ib motoneuron with the corresponding percent of Cpx7A I125M RNA editing. A line of best-fit was generated (black) with 95% confidence intervals displayed (gray). **(D)** Comparison of ADAR transcripts per million (TPM) profile in Is motoneuron with the corresponding percent of Cpx7A I125M RNA editing. **(E)** Quantification of RNA editing percentage for known edit sites in genes encoding the synaptic proteins Synapsin (Syn) and Syntaxin 1A (Syx1A) in comparison to Cpx. Each point represents the percent of editing occurring at the base position of interest in one cell. All cells included for quantification (Ib = 9 cells, Is = 11 cells) contained at least ten reads at all base positions of interest (Syn N15, Syn R19, Syn R20, Cpx I125, and Syx1A M244). **(F)** Representative images of protein loading control (Coomassie blue, top panel) and CK2 phosphorylation ([^32^P] incorporation on autoradiograph, bottom panel) for Cpx7A I125M, N130S (Cpx7A^N130S^) compared to unedited Cpx7A I125, N130 (Cpx7A^N130^) in *in vitro* phosphorylation assays. The absence (-) or presence (+) CK2 in labeling reactions is denoted. **(G)** Quantification of [^32^P] incorporation for the indicated Cpx proteins in *in vitro* CK2 phosphorylation assays. Data are shown as mean ± SEM; \**p* ˂ 0.05, \*\**p* ˂ 0.01, \*\*\**p* ˂ 0.001, \*\*\*\**p* ˂ 0.0001, ns = not significant.

Prior studies proposed editing at adenosine 375 serves to enhance exon-intron base pairing within the 7A pre-spliced mRNA to generate a more favorable double-stranded RNA structure for ADAR to edit the downstream adenosines 388 and 389 that form the AAT codon (N130)^16^. Consistent with this model, RNAseq data from single motoneurons showed editing for both I125 and N130 was far more common than single edits to N130 alone (Figure 4A). Ib neurons expressing Cpx^I125M,N130S^ mRNA were six-fold more abundant that those expressing Cpx^N130S^ alone (I125M, N130S: 9.5 ± 1.5 edit % per cell; N130S: 1.47 ± 0.51 edit % per cell, *p* = 0.0035). A similar ratio was observed in Is neurons (Figure 4A, B). In the 56% of Ib cells expressing Cpx^I125M,N130S^ edited transcripts, an average of 17% of total *cpx* mRNA were of this variant, similar to the 16% of total *cpx* mRNA in the 59% of Is cells that expressed Cpx7A^I125M,N130S^. Rarely, Cpx^I125M,N130S^ represented the only Cpx mRNA detected within a neuron (Figure 4A). In contrast to the more abundant Cpx^I125M,N130S^, only 6% of Ib neurons expressed Cpx^I125M,N130D^ and it represented just 3% of the total *cpx* mRNA in these cells. The Cpx^I125M,N130G^ variant, which requires A-to-I editing at all three adenosines, was observed in 40% of Ib neurons, representing 9% of total *cpx* mRNA in cells in which it was expressed. Similar patterns were observed in Is neurons (Figure 4A, B). For the most abundant edit variant (Cpx^I125M^), no correlation of editing percentage at this site and expression levels of *adar* mRNA in that cell was observed for Ib or Is motoneurons (Figure 4C, D). We conclude that Cpx^I125M,N130S^ is the highest expressed variant in larval motoneurons that alters the N130 residue on the amphipathic helix in Cpx7A.

To determine if RNA editing of other synaptic target genes showed similar stochastic single neuron editing, several additional mRNAs known to undergo RNA editing were examined in the single neuron RNAseq dataset. RNA editing percentages at three sites (N15D, R19G, R20G) within Synapsin (Syn) and one site (M244V) in Syntaxin 1A (Syx1A) were compared in the same motoneurons that edited Cpx7A to I125M (Figure 4E). Both Syn and Syx1A displayed stochastic editing rates that ranged from 0-100% across motoneurons. For example, Syn R20G editing was observed at an average of 55 ± 11.6% (*n* = 11 Is cells), while Syx1A M244V was edited in the same neurons at a rate of only 5.4 ± 3.8% per cell (*n* = 11 Is cells). Similar to Cpx, no significant difference in editing percent of Syn or Syx1A was observed between Ib and Is neurons (Figure 4E). We conclude ADAR-mediated RNA editing is not all-or-none in individual *Drosophila* motoneurons, with stochastic editing at single adenosine base sites and across multiple mRNAs having the potential to generate unique synaptic proteomes within the same population of neurons.

### RNA editing of Cpx7A alters its subcellular localization and functional properties

Given Cpx7A^I125M,N130S^ is the most abundant editing variant on the hydrophilic face of the amphipathic C-terminal helix and resides near the Cpx7B S126 phosphorylation site, the functional significance of RNA editing to N130S or the potential phospho-mimetic version N130D was examined. Unlike Cpx7B, PKA did not phosphorylate unedited Cpx7A^I125,N130^ or Cpx7A^I125M,N130S^ in *in vitro* phosphorylation assays (data not shown). As such, N130S might represent a target for different kinases or instead alter structural properties of the amphipathic helix independent of phosphorylation. Computational analysis of candidate phosphorylation motifs in exon 7A predicted a Casein Kinase 2 (CK2) consensus sequence of S/T E/D at the N130S site that is found in some CK2 targets^40^. To determine if Cpx7A^I125M,N130S^ can be phosphorylated by CK2, *in vitro* phosphorylation assays were performed. Indeed, Cpx7A^I125M,N130S^ was phosphorylated by CK2 (*p* = 0.0003), while unedited Cpx7A^I125,N130^ was not (*p* = 0.9554) (Figure 4F, G), pinpointing N130S as a potential target for CK2 phosphorylation *in vivo*. Although it is unknown if CK2 phosphorylation alters Cpx7A^I125M,N130S^ function, it provides a potential regulatory mechanism downstream of RNA editing.

To examine if RNA editing alters Cpx function at synapses, transgenic UAS rescue lines expressing Cpx7A^I125M,N130S^, Cpx7A^I125M,N130D^ or unedited Cpx7A^I125,N130^ were generated and expressed pan-neuronally using *elav^C155^-*Gal4 in the *cpx^SH1^* null that lacks both Cpx7A and Cpx7B. All three Cpx7A transgenic proteins were overexpressed at similar levels when assayed by western analysis of adult brain lysates (Figure 5A, B; *p* < 0.0001 to control) or anti-Cpx immunostaining at larval muscle 4 NMJs (Figure 5C, D; *p* = 0.0001 to control). Given N130S and N130D alters the hydrophilic face of the C-terminal amphipathic helix that controls SV membrane binding and Cpx localization (Figure 1E), Cpx subcellular distribution at the NMJ was assayed by immunostaining controls (*elav^C155^;; cpx^PE^* (precise excision control for *cpx^SH1^*)), *cpx* nulls (*elav^C155^;; cpx^SH1^*) and the three rescue strains: (1) *elav^C155^;; cpx^SH1^*, UAS-Cpx7A^I125,N130^ (unedited Cpx7A); (2) *elav^C155^;; cpx^SH1^*, UAS-Cpx7A^I125M,N130S^; (3) *elav^C155^;; cpx^SH1^*, UAS-Cpx7A^I125M,N130D^. Endogenous Cpx accumulates along the periphery of presynaptic boutons (Figure 5E), co-localizing with other SV proteins^16^. Lower Cpx levels are found in non-synaptic regions of the axon, resulting in a 3.6-fold synaptic enrichment (Cpx synapse/axon fluorescence ratio) at control NMJs (Figure 5E, F). Expression of unedited Cpx7A^I125,N130^ in the null background resulted in a shift to more Cpx enrichment at synapses, increasing the synapse/axon ratio to 4.7 (Figure 5E, F, *p* = 0.1218). Expression of Cpx7A^I125M,N130S^ or Cpx7A^I125M,N130D^ in the null background had the opposite effect, with a greater fraction of Cpx in non-synaptic regions of the axon (Figure 5E, F). Cpx7A^I125M,N130D^ was enriched 2-fold at synapses, decreasing ~44% compared to controls (*p* = 0.0024) and ~57% compared to unedited Cpx7A^I125,N130^ (*p* ˂ 0.0001). Cpx7A^I125M,N130S^ had a more striking subcellular change, with a nearly one-to-one ratio of its abundance at synapses and along axons (1.16 ± 0.09, *p* ˂ 0.0001 to control). As such, unedited Cpx7A^I125,N130^ is more strongly enriched at synapses, while the predominant splice variant altering the C-terminal amphipathic helix (Cpx7A^I125M,N130S^) distributes equally between axons and synapses. Taken together, these data indicate Cpx distribution at control NMJs likely represents a combinatorial expression of both unedited and edited Cpx proteins.

**Figure 5.**
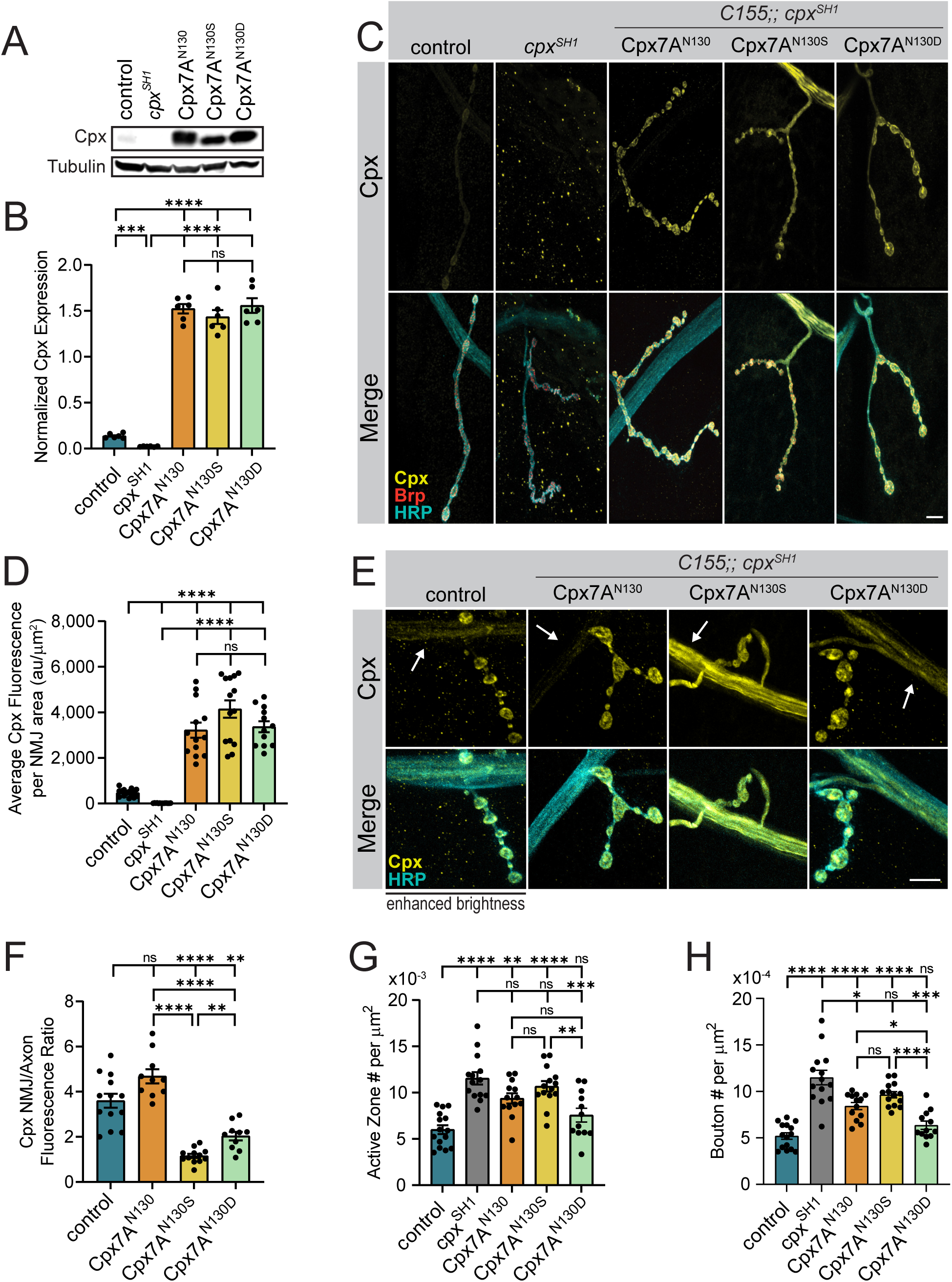
RNA editing of Cpx7A alters its ability to regulate synaptic growth. **(A)** Representative western blot from adult head extracts stained for Cpx or Tubulin (loading control) for the indicated genotypes (control (*elav^C155^-GAL4*;; *cpx^PE^*), *cpx^SH1^* (*elav^C155^-GAL4*;; *cpx^SH1^*), unedited rescue Cpx7A^N130^ (*elav^C155^-GAL4*;; *cpx^SH1^*, UAS-Cpx7A^I125,N130^), Cpx7A^N130S^ rescue (*elav^C155^-GAL4*;; *cpx^SH1^*, UAS-Cpx7A^I125M,N130S^) and Cpx7A^N130D^ rescue (*elav^C155^-GAL4*;; *cpx^SH1^*, UAS-Cpx7A^I125M,N130D^). **(B)** Quantification of Cpx protein levels normalized to Tubulin from western blots of the indicated genotypes. **(C)** Immunostaining of 3^rd^ instar larval muscle 4 NMJs at segment A3 for the indicated genotypes with antibodies against Cpx (yellow, upper panels), Brp (red) and anti-HRP (cyan). Scale bar = 10 µm. **(D)** Quantification of total Cpx fluorescence within the HRP-positive area at muscle 4 NMJs for the indicated genotypes. **(E)** Representative staining of 3^rd^ instar muscle 4 NMJs and axons of the indicated genotypes from segment A3 with antibodies against Cpx (yellow) and HRP (cyan). Cpx staining in axons is denoted with white arrows. The brightness of anti-Cpx staining was enhanced in controls to demonstrate the lower amounts of Cpx normally found in non-synaptic regions of the axon. Scale bar = 10 µm. **(F)** Quantification of the Cpx NMJ/axon fluorescent ratio for the indicated genotypes. **(G)** Quantification of mean AZ number per muscle area at muscle 4 NMJs of the indicated genotypes. **(H)** Quantification of mean synaptic bouton number per muscle area at muscle 4 NMJs of the indicated genotypes. Data are shown as mean ± SEM; \**p* ˂ 0.05, \*\**p* ˂ 0.01, \*\*\**p* ˂ 0.001, \*\*\*\**p* ˂ 0.0001, ns = not significant.

To directly test the functional impacts of RNA editing of Cpx *in vivo*, morphological and electrophysiological assays were performed in rescue lines. Quantification of synaptic bouton and AZ number revealed significant differences in the ability of different Cpx7A proteins to rescue synaptic overgrowth in *cpx^SH1^* null larvae (Figure 5G, H). Cpx7A^I125M,N130D^ fully rescued the increased number of AZs (*p* = 0.3658, Figure 5G) and boutons (*p* = 0.3722, Figure 5H), returning synaptic morphology to control levels. In contrast, Cpx7A^I125M,N130S^ displayed the weakest rescue, with ~80% more AZs and boutons than controls (*p* ˂ 0.0001), slightly less than the doubling of AZ and bouton number observed in *cpx^SH1^* (Figure 5G, H). Expression of unedited Cpx7A^I125,N130^ in the null background resulted in a partial rescue of the morphological defects (AZ#: *p* = 0.0013, bouton#: *p* < 0.0001). Electrophysiological analysis of spontaneous fusion rates revealed a similar pattern of rescue as observed for synaptic growth (Figure 6A, B), consistent with enhanced mini frequency being the primary driver for synaptic over-proliferation in *cpx* mutants. Like the full rescue of synaptic overgrowth, Cpx7A^I125M,N130D^ returned mini frequency to control levels (*p* > 0.9999). In contrast, Cpx7A^I125M,N130S^ expression failed to properly clamp spontaneous release, similar to its inability to rescue synaptic overgrowth. Compared to the 102 Hz mini frequency at *cpx^SH1^* null NMJs, Cpx7A^I125M,N130S^ expression was able to reduce spontaneous fusion by only ~50% to 47 Hz (*p* < 0.0001 to control). Rescue with unedited Cpx7A^I125,N130^ created a more effective fusion clamp, decreasing mini frequency to 13 Hz (*p* = 0.0034 to control), although this rate was still ~3-fold higher than controls that also contain endogenous Cpx7B or the Cpx7A^I125M,N130D^ rescue. RNA editing of Cpx7A also impacted evoked release, though not as severely as spontaneous release. Both unedited and edited Cpx7A improved eEJC amplitude and release kinetics compared to null mutants (Figure 6C-F). Similar to spontaneous release, Cpx7A^I125M,N130D^ showed the strongest rescue for evoked responses (*p* = 0.0371 to control), while Cpx7A^I125M,N130S^ and Cpx7A^I125M,N130^ displayed intermediate rescues. We conclude that RNA editing of Cpx7A regulates its functional properties, with N130D and N130S displaying several opposing effects compared to unedited Cpx7A. The N130D edit improved Cpx’s ability to clamp spontaneous release, in contrast to the N130S edit which reduced clamping and failed to prevent synaptic overgrowth. Based on the distinct phenotypes of N130S and N130D, it seems unlikely the primary effect of N130S is downstream of phosphorylation, but instead reflects a change to the functional properties of the C-terminal amphipathic helix. Alternatively, the N130D change does not act as a phospho-mimetic and instead alters the Cpx7A C-terminal helix in a distinct manner.

**Figure 6.**
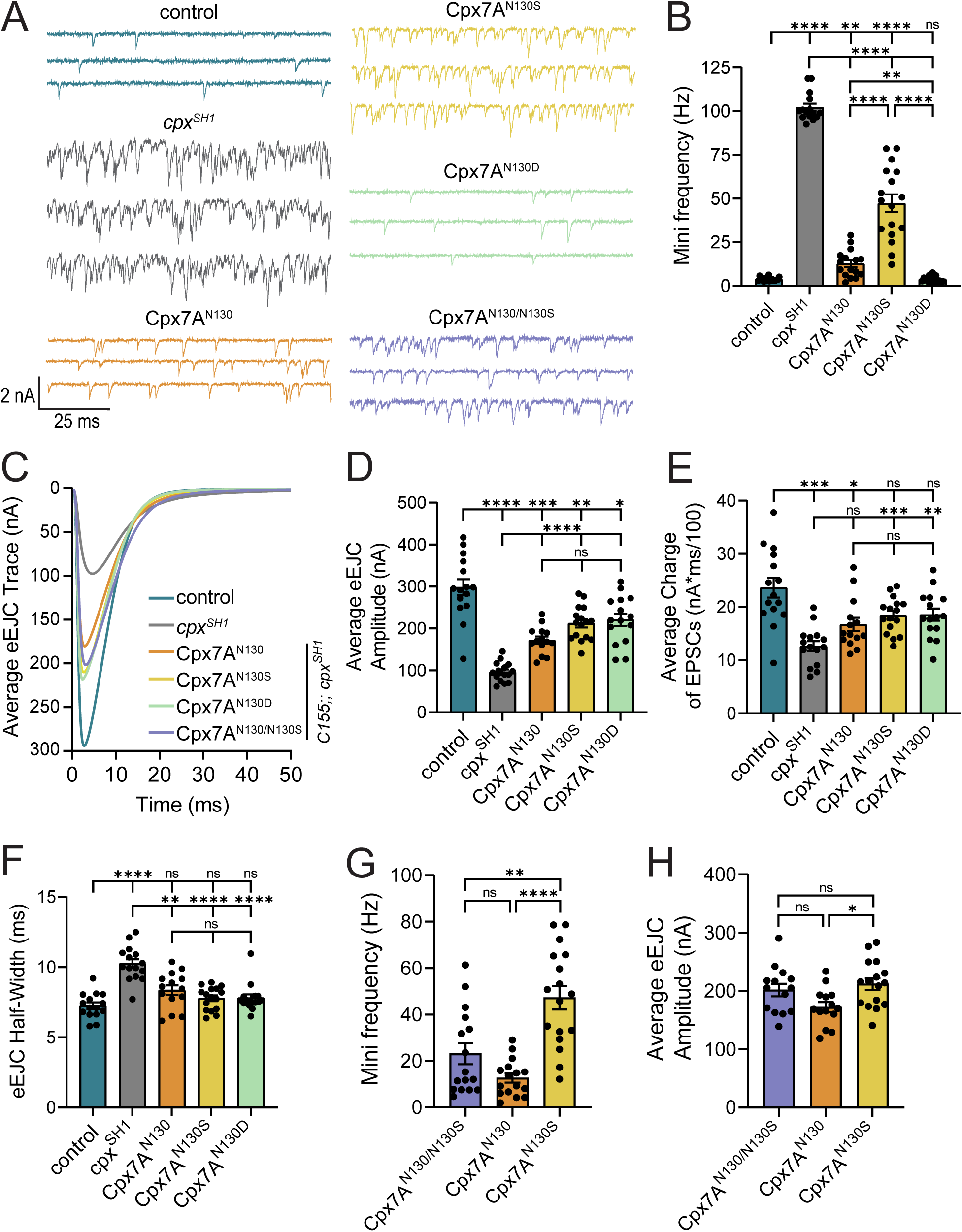
RNA editing of Cpx7A alters it ability to regulat neurotransmitter release. **(A)** Representative postsynaptic current recordings of spontaneous release at muscle 6 NMJs in control (blue, *elav^C155^-GAL4*;; *cpx^PE^*), *cpx^SH1^* (gray, *elav^C155^-GAL4*;; *cpx^SH1^*), unedited Cpx7A^N130^ rescue (orange, *elav^C155^-GAL4*;; *cpx^SH1^*, UAS-Cpx7A^I125,N130^), Cpx7A^N130S^ rescue (yellow, *elav^C155^-GAL4*;; *cpx^SH1^*, UAS-Cpx7A^I125M,N130S^), Cpx7A^N130D^ rescue (green, *elav^C155^-GAL4*;; *cpx^SH1^*, UAS-Cpx7A^I125M,N130D^) and co-expression of Cpx7A^N130/N130S^ rescue (purple, *elav^C155^-GAL4*;; *cpx^SH1^*, UAS-Cpx7A^I125,N130^/*cpx^SH1^*, UAS-Cpx7A^I125M,N130S^). **(B)** Quantification of average spontaneous release rate for the indicated genotypes. **(C)** Average traces of evoked EJC responses for the indicated genotypes. **(D)** Quantification of average eEJC amplitude for the indicated genotypes. **(E)** Quantification of average evoked release charge obtained by measuring total release over time following single action potentials. **(F)** Average evoked EJC half-width change for each genotype. **(G)** Quantification of average spontaneous release rate for co-expression of Cpx7A^N130/N130S^ rescue compared to Cpx7A^N130^ rescue and Cpx7A^N130S^ rescue alone in the *cpx^SH1^* mutant background. **(H)** Quantification of average eEJC amplitude for co-expression of Cpx7A^N130/N130S^ rescue compared to Cpx7A^N130^ rescue and Cpx7A^N130S^ rescue alone in the *cpx^SH1^* mutant background. All recordings were performed in 2.0 mM external Ca^2+^ saline. Data are shown as mean ± SEM; \**p* ˂ 0.05, \*\**p* ˂ 0.01, \*\*\**p* ˂ 0.001, \*\*\*\**p* ˂ 0.0001, ns = not significant.

RNAseq analysis of single neuron editing demonstrated most motoneurons express a combination of edited Cpx7A proteins (Figure 4A, B), with Cpx7A^I125M,N130S^ representing the most abundant edit to the N130 residue. To determine if co-expression of unedited Cpx7A^I125,N130^ with Cpx7A^I125M,N130S^, as would be observed *in vivo*, results in competition for Cpx interactions with the SNARE fusion machinery, the two proteins were co-expressed in the *cpx^SH1^* null background. If unedited Cpx7A restored mini frequency to baseline rates, RNA editing would likely have effects on synaptic transmission only when neurons predominantly express edited Cpxs. Alternatively, if co-expression of Cpx7A^I125M,N130S^ increased mini frequency beyond unedited Cpx7A^I125,N130^ rescues alone, a combinatorial role for unique Cpx edit variants to modulate presynaptic output from a neuron would be more likely. An intermediate effect on spontaneous fusion was observed when unedited Cpx7A^I125,N130^ and Cpx7A^I125M,N130S^ were co-expressed in *cpx^SH1^*. Co-expression of the two proteins resulted in a spontaneous release rate of 23 Hz (Figure 6A, G), distinct from the 47 Hz in Cpx7A^I125M,N130S^ (*p* = 0.0036). Co-expression of unedited Cpx7A^I125,N130^ and Cpx7A^I125M,N130S^ also displayed a larger evoked response than unedited Cpx7A^I125,N130^ alone, suggesting the N130S acts in a dominant fashion for promoting evoked release (Figure 6C, H). These data suggest co-expression of different Cpx edit variants can independently interface with the SV release machinery, consistent with stochastic RNA editing of Cpx acting as a mechanism for generating release heterogeneity across individual neurons.

## Discussion

Mechanisms controlling SNARE complex assembly dynamics provide attractive sites for regulatory control of presynaptic neurotransmitter release. Several SNARE binding proteins act at multiple steps of the SV cycle to chaperone SNARE proteins and regulate their ability to zipper together to form the four-stranded alpha-helical bundle that drives fusion^1,41^. Cpx and Syt1 play key roles at a late stage of SNARE assembly to control whether SVs fuse through the evoked or spontaneous release pathway. The Cpx C-terminal amphipathic helix has emerged as an important site for regulatory control of the protein^15^. In addition to acting as a membrane curvature sensor that can tether Cpx to SVs^24–26,42,43^, the C-terminal domain can clamp fusion by blocking SNARE assembly^44^, remodel membranes to regulate fusion pore dynamics^45^, directly stimulate SNARE-mediated assembly^46^, or compete with Syt1 for membrane binding^47^. In addition, post-translational modifications to this domain can alter Cpx function^11,12,46,48–50^, defining presynaptic plasticity mechanisms that directly impinge on SNARE-mediated fusion.

In the current study, we examined the functional consequences of alternative splicing and RNA editing to the C-terminus of *Drosophila* Cpx. Endogenous expression of either Cpx7A or Cpx7B was sufficient to prevent synaptic overgrowth and the dramatic elevation of spontaneous release rates observed in null mutants. Indeed, across all manipulations of Cpx splicing and RNA editing, large increases in mini frequency were always associated with over proliferation of synapses. These observations support the linkage between spontaneous fusion and retrograde signaling mechanisms that drive one form of activity-dependent synaptic growth at *Drosophila* NMJs^11,51^. Endogenously expressed Cpx7A or Cpx7B alone were not sufficient to fully recapitulate evoked release observed in controls, suggesting both are required *in vivo*. Given differences in expression level between Cpx7A and Cpx7B, it was surprising that endogenous Cpx7B could support Cpx function at larval NMJs. Loss of the predominant Cpx7A isoform in the *cpx^Δ7A^* splicing mutant reduced overall Cpx levels by 87% in adult heads and 85% at NMJs. In contrast, loss of Cpx7B in *cpx^Δ7B^* reduced total Cpx levels by only 10% in adult heads and did not cause a detectable decrease at NMJs. We considered the possibility that a truncated Cpx might still be produced that lacked the 7A exon in *cpx^Δ7A^* mutants that supported some release on its own. However, western analysis showed that only traces of a truncated protein as a much fainter lower molecular weight band that was barely detectable in *cpx^Δ7A^*, dramatically less than the already reduced levels of Cpx7B. These data indicate loss of the C-terminal domain destabilizes Cpx and leads to its degradation, similar to our prior observations on a *Drosophila* Cpx truncating mutant (*cpx^572^*) containing a stop codon at the end of exon 6^16^.

Given Cpx7B clamped spontaneous fusion and promoted evoked release at only ~15% of the Cpx7A expression level, the two splice variants likely have intrinsic differences in their activity. Like mammalian CPX1 and CPX2 that are present at most central nervous system (CNS) synapses, Cpx7B lacks the C-terminal CAAX motif, potentially endowing these non-prenylated Cpx proteins with greater mobility to interface with Syt1 and SNAREs and enhance release kinetics. Indeed, the onset of evoked release was slightly faster in larvae expressing only Cpx7B. While Cpx7B was able to reduce elevated mini frequency 28-fold compared to nulls, it was only able to rescue ~50% of evoked release, suggesting Cpx abundance is more critical for promoting evoked fusion than clamping minis. The estimated requirements for zippering of 3 to 11 SNARE complexes in a radial assembly for a single SV to undergo action potential-triggered fusion^52–55^ provides a candidate mechanism for Cpx expression to differentially impact these two release pathways. For spontaneous release, a smaller number of Cpx proteins could block enough SNARE zippering events to prevent reaching the minimum required for fusion. For rapid Ca^2+^-triggered release, excess Cpxs would be needed to bind more SNARE complexes in the radial assembly at the SV-plasma membrane interface to control Syt1 activity and SNARE zippering.

We previously discovered that PKA phosphorylation of S126 in the Cpx7B C-terminus enhances spontaneous release and promotes structural and functional synaptic plasticity at larval NMJs^11^. PKA phosphorylation of S126 also controls fusion pore dilation and cargo release from *Drosophila* dense core vesicles^50^, indicating changes to the C-terminal amphipathic domain of Cpx modulate its function in membrane fusion. Although Cpx7A lacks this PKA phosphorylation site, it undergoes RNA editing to generate up to eight unique C-terminal sequence variants (I125 with N130, S130, D130, or G130 and M125 with N130, S130, D130 or G130). Single-cell RNAseq revealed multiple Cpx7A editing variants are simultaneously expressed in Ib and Is motoneurons. As such, ADAR-mediated RNA editing does not act in an “all-or-none” fashion in *Drosophila* motoneurons, but instead stochastically deaminates A-to-I residues with distinct efficiencies. Generally, unedited Cpx7A was the dominant form of Cpx in motoneurons, with editing variants represented at lower levels. However, in some cases, a single edit variant was the only Cpx mRNA expressed in that neuron (Figure 4A). The amount of Cpx editing was not correlated with ADAR expression, similar to the dissociation of editing percentage and ADAR abundance in pooled RNAseq data from adult *Drosophila* neurons over a broader range of RNA editing targets^56^. Although the percentage of Cpx editing variants were similarly expressed in the Ib and Is motoneurons, analysis of FACS-sorted neuronal populations from adult *Drosophila* brains revealed RNA editing of Cpx7A was even more robust and diverse in these neurons, with 22% of *cpx* mRNAs encoding N130S and 23% encoding N130G^56^. As *Drosophila* ADAR itself is subject to developmentally-regulated auto-editing that can change its enzymatic activity^57^, the frequency of Cpx editing might be modulated by intrinsic activity, allowing more dynamic changes to Cpx function within single neurons.

To determine if Cpx7A edit variants within a single neuron alter presynaptic output, transgenic rescues were used to assay their function. The most prominent variant Cpx7A^I125M,N130S^ displayed altered synaptic distribution, with more of the protein observed in axons. It also failed to clamp spontaneous fusion, resulting in synaptic overgrowth. We found that Cpx7A^I125M,N130S^ could be phosphorylated by CK2 *in vitro*. CK2 phosphorylates multiple synaptic proteins, including Syx1A^58,59^, Syt1^60^ and mammalian CPX1^48^. Phosphorylation of the C-terminus of mammalian CPX1 by CK2 alters its SNARE-binding affinity^48^, while mutation of the CK2 phosphorylation site (S115) prevents CPX1 from stimulating liposomal fusion^46^. In *Drosophila*, presynaptic CK2 has been demonstrated to control synaptic stability by regulating Ankyrin2 function^40^, preventing a detailed analysis of its role in neurotransmitter release due to synapse loss. Although CK2 can phosphorylate Cpx7A^I125M,N130S^ *in vitro*, it remains unclear if this is relevant to Cpx function *in vivo*. In particular, the putative phospho-mimetic variant Cpx7A^I125M,N130D^ was able to fully clamp spontaneous release and support normal synaptic growth, outperforming even unedited Cpx7A. As such, Cpx7A^I125M,N130S^ may instead alter the structure or binding properties of the amphipathic helix in unique ways to prevent normal interactions with SNAREs or other protein/lipid targets.

Given the stochastic expression of Cpx edited proteins within single motoneurons, and the requirement for multiple SNARE complexes to drive fusion, we assayed if co-expression of Cpx7A^I125M,N130S^ with unedited Cpx7A^I125,N130^ could allow distinct Cpx proteins to independently alter release output. Indeed, the N130S isoform acted in a partially dominant manner, preventing unedited Cpx7A from fully clamping spontaneous fusion, while supporting higher levels of evoked release. These data support a model where each Cpx variant has independent access to assembling SNAREs, allowing each to fine-tune presynaptic output in unique ways. Indeed, disruptions of the amphipathic helix in *C. elegans* Cpx alter SV release^24,43^, further highlighting a key role for this domain.

Beyond stochastic RNA editing of Cpx7A, we observed similar heterogeneity in the percent of RNA editing in Synapsin and Syx1A mRNAs across *Drosophila* motoneurons. As such, stochastic RNA editing across multiple mRNAs is likely to generate unique synaptic proteomes within the same neuronal population and contribute to heterogenous properties of individual cells with similar transcriptomes. Such a mechanism would be a robust way to alter multiple features of neuronal output given ADAR editing alters the function of proteins that contribute to both synaptic release and membrane excitability^19,61–64^.

## Methods

### Drosophila stocks

Flies were cultured on standard medium and maintained at 25 °C. Late 3^rd^ instar larvae were used for imaging and electrophysiological experiments. Western blots were performed on adult brain extracts. Males were used for experiments unless otherwise noted. Experiments were performed in a *w*^1118^ (Bloomington *Drosophila* Stock Center #3605) genetic background unless otherwise noted.

### Transgenic constructs

QuikChange Lightning (Agilent) was used for site-directed mutagenesis on unedited Cpx7A to generate specified *Cpx* edit variants that were subcloned into modified pValum construct, as previously described^11^. The resulting constructs were injected into a yv;;attP 3^rd^ chromosome docking strain by BestGene Inc. (Chino Hills, CA, USA). UAS lines were recombined into the *cpx^SH1^* null mutant background and *elav^C155^*-GAL4 (BDSC #8765) was used for pan-neuronal expression of transgenes.

### Generation of CRISPR-modified Cpx strains

Two endogenous *Cpx* truncation lines were generated (*cpx^Δ7A^* and *cpx^Δ7B^*) using a CRISPR genome engineering approach. Four guide RNAs (gRNAs) flanking the splice acceptor site of exon 7A or 7B were selected using the CRISPR Optimal Target Finder^65^. gRNAs were cloned into the pCFD5 expression vector (Addgene #73914)^66^ and donor constructs were generated to encode a floxed P3>DsRed reporter cassette (Addgene #51434) in the reverse orientation flanked with one kb homology arms upstream and downstream of the splice acceptor site of either exon 7A or 7B by Gibson assembly protocol using NEBuilder HighFidelity DNA Assembly Cloning Kit (E5520). An early stop codon was inserted between homology arms for each respective exon construct, with several amino acid coding sequences maintained to preserve proper exon splicing. gRNA binding sites of donor template were mutated using silent mutations that did not alter amino acid sequence. Template and gRNA plasmids were co-injected into *vasa*-Cas9 embryos (BDSC #56552) by BestGene Inc (Chino Hills, CA, USA) and Ds>Red positive transformants were selected by BestGene Inc (Chino Hills, CA, USA). The modified locus with stop codons inserted into exon 7A or 7B were confirmed by sequencing.

### Locomotion analysis

Adult geotaxis was measured in adult male flies aged 2-3 days as previously described^67^. Briefly, eight cohorts of ten adult males (80 total flies) per genotype were separated after eclosion and allowed to recover from CO_2_ for 24 hours on standard fly medium. After 24 hours, each cohort was moved to a chamber made from two clear plastic vials taped together, with a line drawn around the lower vial 8 cm from the bottom. Each cohort was allowed to acclimate in the chamber for five minutes before assays began. Negative geotaxis was measured as percent of the cohort that crossed the 8 cm line within ten seconds after being tapped to the bottom of the chamber. Each cohort was subjected to ten rounds of negative geotaxis assay, with a one-minute rest period between each. The percent of flies that crossed the 8 cm line after each round was averaged to produce a pass rate per cohort. Larval crawling was assayed in 3^rd^ instar larvae of both sexes, as previously described^68,69^. Larvae were briefly washed in room-temperature water before placing onto the center of a 5 cm petri dish containing 2% agarose, with five animals from a single genotype placed together (*n* = 10 larvae per genotype). The petri dish was placed over a grid and velocity was measured as average distance (in mm) traveled during the first 30 seconds following placement.

### Immunohistochemistry

Larvae were dissected in hemolymph-like HL3.1 solution (in mM: 70 NaCl, 5 KCl, 4 MgCl_2_, 10 NaHCO3, 5 trehalose, 115 sucrose, 5 HEPES, pH 7.2) and fixed in 4% paraformaldehyde for 18 minutes. Larvae were washed three times for five minutes with PBST (PBS containing 0.1% Triton X-100), followed by a thirty-minute incubation in block solution (5% NGS in PBST). Fresh block solution and primary antibodies were then added. Samples were incubated overnight at 4°C and washed with two short washes and three extended 20 minutes washed in PBST. PBST was replaced with block solution and fluorophore-conjugated secondary antibodies were added. Samples were incubated at room temperature for two hours. Finally, larvae were rewashed with PBST and mounted in Vectashield (Vector Laboratories, Burlingame, CA). Antibodies used for this study include: mouse anti-Brp, 1:500 (NC82; Developmental Studies Hybridoma Bank (DSHB), Iowa City, IA)); rabbit anti-Cpx, 1:5000^7^; goat anti-rabbit Alexa Fluor 488, 1:500 (A-11008; ThermoFisher Scientific, Waltham, MA, USA); goat anti-mouse Alexa Fluor 546, 1:500 (A-11030; ThermoFisher); DyLight 649 conjugated anti-HRP, 1:500 (#123-605-021; Jackson Immuno Research, West Grove, PA, USA).

### Confocal imaging and imaging data analysis

Imaging was performed on a Zeiss Pascal confocal microscope (Carl Zeiss Microscopy, Jena, Germany) using a 63X 1.3 NA oil-immersion objective (Carl Zeiss Microscopy). Images were processed with the Zen (Zeiss) software. A 3D image stack was acquired for each NMJ imaged (muscle 4 Ib NMJ of abdominal segment A3) and merged into a single plane for 2D analysis using FIJI image analysis software^70^. No more than two NMJs were analyzed per larva. Anti-HRP labeling was used to identify neuronal anatomy (axons and NMJs) and quantify synaptic bouton number and NMJ area. Brp puncta quantification was used to measure AZ number. Muscle 4 area was used to normalize quantifications for muscle surface area. For Cpx fluorescence quantification, the HRP-positive area was used to outline NMJs and axons. Total Cpx fluorescent intensity was measured in the outlined area, with background fluorescence of mean pixel intensity of non-HRP areas subtracted. For NMJ/axon ratios, background subtracted mean NMJ Cpx fluorescence was compared to background subtracted mean axon Cpx fluorescence within the same image.

### Two-electrode voltage-clamp electrophysiology

Postsynaptic currents were recorded from 3^rd^ instar muscle 6 at segment A3 using two-electrode voltage clamp with a −80 mV holding potential. Experiments were performed in room temperature HL3.1 saline solution (in mM, 70 NaCl, 5 KCl, 10 NaHCO3, 4 MgCl2, 5 trehalose, 115 sucrose, 5 HEPES, pH 7.2). Final [Ca^2+^] was adjusted to 2 mM unless otherwise noted. Motor axon bundles were cut and suctioned into a glass electrode and action potentials were stimulated at 0.5 Hz (unless indicated) using a programmable stimulator (Master8, AMPI; Jerusalem, Israel). Data acquisition and analysis was performed using Axoscope 10.0 and Clampfit 10.0 software (Molecular Devices, Sunnyvale, CA, USA) and inward currents were labeled on a reverse axis for clarity.

### Western blot analysis

Western blotting of adult head lysates (three heads per sample with one head loaded per lane) was performed using standard laboratory procedures with mouse anti-Tubulin (T5168; Sigma) at 1:10000 (UAS rescue experiments) or 1:1000000 (CRISPR experiments) and rabbit anti-Cpx at 1:5000. IRDye 680LT-conjugated goat anti-mouse, 1:5000 (926–68020; LICOR) and IRDye 800CW conjugated goat anti-rabbit, 1:5000 (926-32211; LICOR) were used as secondary antibodies. Blocking was performed in a solution containing four parts TBS (10 mM Tris Base pH 7.5, 150 mM NaCl) to one part Blocking Buffer (Rockland) for one hour. Antibody incubations were performed in a solution containing four parts TBST (1X TBS with 1% Tween-20) to one part Blocking Buffer. A LI-COR Odyssey Imaging System (LI-COR Biosciences, Lincoln, MA, USA) was used for visualization and analysis was performed using FIJI image analysis software. Relative Cpx expression was calculated by normalizing to Tubulin intensity.

### Purification of Complexin for in vitro phosphorylation assay*s*

QuikChange Lightning (Agilent) was used for site-directed mutagenesis of unedited Cpx7A to generate Cpx7A^I125M,N130S^ (termed N130S). Recombinant Cpx fused with GST was expressed in E. coli (BL21) and purified using glutathione sepharose 4B (Fisher Scientific). Peak fractions were concentrated and further purified by gel filtration as previously described^11^. *In vitro* kinase assays were performed using purified recombinant Cpx proteins and the catalytic subunit of CK2 (C70-10G, SignalChem). Briefly, 10 mg of purified GST-fusion protein (unedited Cpx7A^I125,N130^ or edited Cpx7A^I125M,N130S^) was used per reaction and incubated with 2,500 units of recombinant kinase and [^32^P]ATP (Perkin Elmer). Reaction products were separated by SDS-PAGE and gels were stained with Bio-Safe Coomassie Blue (Bio-Rad), dried, and exposed to autoradiography film at room temperature. Mean integrated density of each band was quantified using FIJI and relative density of phospho-Cpx (pCpx) was calculated by normalizing to input band intensity determined by Coomassie staining.

### RNAseq analysis of RNA editing

RNAseq data from 105 single MN1-Ib and 101 single MNISN-Is 3^rd^ instar larval motoneurons obtained using isoform Patchseq protocols^33^ were analyzed using the Integrative Genomics Viewer (IGV)^71^. To create single-cell Cpx RNA editing expression profiles, single RNA reads were analyzed for Cpx and included in the analysis if all three C-terminal Cpx7A edited bases were represented on a continuous single read. The percent of each Cpx7A edit variant was determined by the number of edited variant reads divided by total RNA reads for each cell, creating an RNA editing profile for each neuron. To compare single base editing across different genes, the edit percent at each base of interest was analyzed and compared to known edits in other genes within the same neurons. Neurons were excluded if each base of interest did not contain ten or more reads for all edits of interest.

### Experimental design and statistical analysis

Statistical analysis and plot generation was performed using GraphPad Prism software version 9.5.1 (San Diego, CA, USA). Appropriate sample size was determined using a normality test. Statistical significance for comparisons of two groups was determined by a two-tailed Student’s t-test. For comparisons of three or more groups of data, a one-way ANOVA followed by Tukey’s Multiple Comparisons test was used to determine significance when the largest sample standard deviation was no more than twice as large as the smallest sample standard deviation. When the largest sample standard deviation was more than twice as large as the smallest sample standard deviation, a Brown-Forsythe and Welch’s one-way ANOVA followed by Dunnett’s T3 Multiple Comparisons test was used to determine significance. For comparisons of two factors with three or more groups of data, as described in Figure 4A, G, a two-way ANOVA followed by Tukey’s Multiple Comparisons test was used. For **Figure 4C-D**, a line of best-fit was generated with 95% confidence intervals displayed. The mean of each distribution is plotted in figures with individual datapoints also shown. Error bars represent ± SEM. Asterisks indicate the following *p*-values: *, *p*˂0.05; **, *p*˂0.01; ***, *p*˂0.001; ****, *p*˂0.0001, with ns = not significant.

## Acknowledgements

We thank the Bloomington *Drosophila* Stock Center (Indiana University, Bloomington, IN; NIH P40OD018537), the Developmental Studies Hybridoma Bank (University of Iowa, Iowa City, IA), Dina Volfson and Richard Cho for assistance with experimental methods and *Drosophila* strains, and members of the Littleton lab for helpful discussions. This work was supported by The JPB Foundation and a National Institutes of Health grant (NS40296) to J.T.L., and a NSF grant (1122374) to E.A.B.

## Author contributions

E.A.B: Conceptualization, Methodology, Formal Analysis, Investigation, Writing – Original Draft, Visualization; Z.G.: Formal Analysis, Investigation; S.K.J: Resources; J.T.L: Conceptualization, Methodology, Writing – Original Draft, Supervision, Funding Acquisition.

